# Evidence for a Nuclear Role for *Drosophila* Dlg as a Regulator of the NURF Complex

**DOI:** 10.1101/2021.04.18.440284

**Authors:** Katherine A. Sharp, Mark J. Khoury, Frederick Wirtz-Peitz, David Bilder

**Affiliations:** Department of Molecular and Cell Biology, University of California-Berkeley, Berkeley CA 94720; Department of Genetics, Harvard Medical School, Boston, MA

## Abstract

Scrib, Dlg, and Lgl are basolateral regulators of epithelial polarity and tumor suppressors whose molecular mechanisms of action remain unclear. We used proximity biotinylation to identify proteins localized near Dlg in the *Drosophila* wing imaginal disc epithelium. In addition to expected membrane- and cytoskeleton-associated protein classes, nuclear proteins were prevalent in the resulting mass spectrometry data set, including all four members of the NURF chromatin remodeling complex. Subcellular fractionation demonstrated a nuclear pool of Dlg and proximity ligation confirmed its position near the NURF complex. Genetic analysis showed that NURF activity is also required for the overgrowth of *dlg* tumors, and this growth suppression correlated with a reduction in Hippo pathway gene expression. Together, these data suggest a nuclear role for Dlg in regulating chromatin and transcription through a more direct mechanism than previously thought.

**Highlight Summary:** Proximity proteomics is used as an entry point towards identifying partners of the polarity-regulating tumor suppressor Dlg. A nuclear pool of the protein associated with NURF chromatin remodelers is revealed, along with evidence of functional interactions during growth regulation.

## Introduction

Discs-large (Dlg), Scribble (Scrib), and Lethal giant larvae (Lgl) are evolutionarily conserved polarity-regulating proteins found at the basolateral membranes of epithelial cells, where they restrict the localization of the aPKC and Par complexes to the apical region of the cell (Elsum *et al*., 2012; Rodriguez-Boulan and Macara, 2014; Campanale *et al*., 2017). They can also regulate the formation and maintenance of cell junctions, the division axis of epithelial cells, and the asymmetric division of stem cells (Woods *et al*., 1996; Tepass and Tanentzapf, 2001; Albertson and Doe, 2003; Bilder *et al*., 2003; Johnston *et al*., 2009; Bergstralh *et al*., 2013; Rodriguez-Boulan and Macara, 2014; Campanale *et al*., 2017; Nakajima *et al*., 2019). In *Drosophila* epithelia, loss of any one of these three proteins causes not only loss of polarity but also neoplastic transformation and tumorous overgrowth (Bilder *et al*., 2000; Bilder, 2004; Hariharan and Bilder, 2006; Humbert *et al*., 2008). Overgrowth results from an aberrant transcriptional program that is driven by Yorkie (Yki), the transcriptional activator of the Hippo pathway (Hariharan and Bilder, 2006; Grzeschik *et al*., 2010; Zhu *et al*., 2010; Doggett *et al*., 2011; Sun and Irvine, 2011; Bunker *et al*., 2015).

Dlg, Scrib, and Lgl are conserved in vertebrates where they each have multiple homologs (Elsum *et al*., 2012). As in flies, the vertebrate proteins have been implicated in regulation of tumor growth, with changes detected in a variety of human cancers (Halaoui and McCaffrey, 2015). They are also involved in apicobasal polarity and formation of both adherens and occluding tight junctions in epithelial cells (Su *et al*., 2012; Choi *et al*., 2019) and have a similar role in endothelial cells (Elsum *et al*., 2012; Lizama and Zovein, 2013; Worzfeld and Schwaninger, 2016). Further, Dlg homologs regulate the migration of epithelial cells during development. Mutations in these genes can result in cleft palate (Caruana and Bernstein, 2001), hydrocephalus (Nechiporuk *et al*., 2007), and defects in renal and urogenital systems (Iizuka-Kogo *et al*., 2007; Nechiporuk *et al*., 2007; Elsum *et al*., 2012).

Despite the importance of Scrib, Dlg, and Lgl for development, homeostasis, and disease, we still have a limited understanding of their molecular functions and the mechanisms by which they regulate cell biology and gene expression. Scrib and Dlg are multivalent “scaffolding” proteins containing a variety of protein-protein interaction domains and motifs. Scrib contains 4 PDZ domains and a leucine-rich repeat domain, while Dlg contains three PDZ domains, one SH3 domain, and a catalytically-dead guanylate kinase domain (Tepass and Tanentzapf, 2001; Elsum *et al*., 2012; Su *et al*., 2012; Campanale *et al*., 2017). Understanding the function of these scaffolds will require defining the proteins with which they interact as well as how those interactions change and are regulated over space and time within cells. However, PDZ and other domains are thought to facilitate weak and transient interactions, often involving plasma membrane-embedded receptors, making it difficult to define the full complement of proteins with which they interact using traditional biochemical methods (Amacher *et al*., 2020). Numerous prior efforts have used co-immunopreciptitation with mass spectrometry (IP-MS) to identify binding partners for Dlg, Scrib, and Lgl (Audebert *et al*., 2004; Van Campenhout *et al*., 2011; Anastas *et al*., 2012; Belotti *et al*., 2013; Michaelis *et al*., 2013; Nagasaka *et al*., 2013; Ivarsson *et al*., 2014; Waaijers *et al*., 2016; Drew *et al*., 2017; Dash *et al*., 2018; Portela *et al*., 2018; Nakajima *et al*., 2019) (reviewed in (Stephens *et al*., 2018)). However, such experiments have yielded largely non-overlapping lists of binding partners and relatively few mechanistic insights. The limited utility of some IP-MS approaches may additionally derive from the use of non-epithelial cell lines in which functionally important interactions may not exist.

We sought a different approach that could be carried out in intact epithelial cells and that would not rely on strong, stable interactions between proteins. We therefore turned to proximity-based biotin labeling using the APEX2 enzyme. APEX2 is an ascorbate-peroxidase derived from the pea plant that, in the presence of hydrogen peroxide, catalyzes the conversion of phenol into a phenoxyl radical. In cells supplied with biotin-phenol as a substrate, APEX2 covalently labels proteins with biotin within a 20nm radius of the enzyme (Figure 1A). By leveraging the strength and specificity of streptavidin-biotin binding, labeled proteins can then be efficiently and cleanly isolated and identified by MS (Martell *et al*., 2012; Rhee *et al*., 2013; Hung *et al*., 2014, 2016; Lam *et al*., 2014; Chen *et al*., 2015). Proximity-based biotinylation has been used in a variety of experimental systems to identify catalogs of proteins associated with a particular organelle or localized to a particular subcellular region. Importantly, this method can also capture protein-protein interactions that cannot be isolated by more conventional methods (Rhee *et al*., 2013; Van Itallie *et al*., 2013; Hung *et al*., 2014; Chen *et al*., 2015; Gingras *et al*., 2019; Mannix *et al*., 2019; Trinkle-Mulcahy, 2019; Bagci *et al*., 2020; Tan *et al*., 2020).

**Figure 1:**
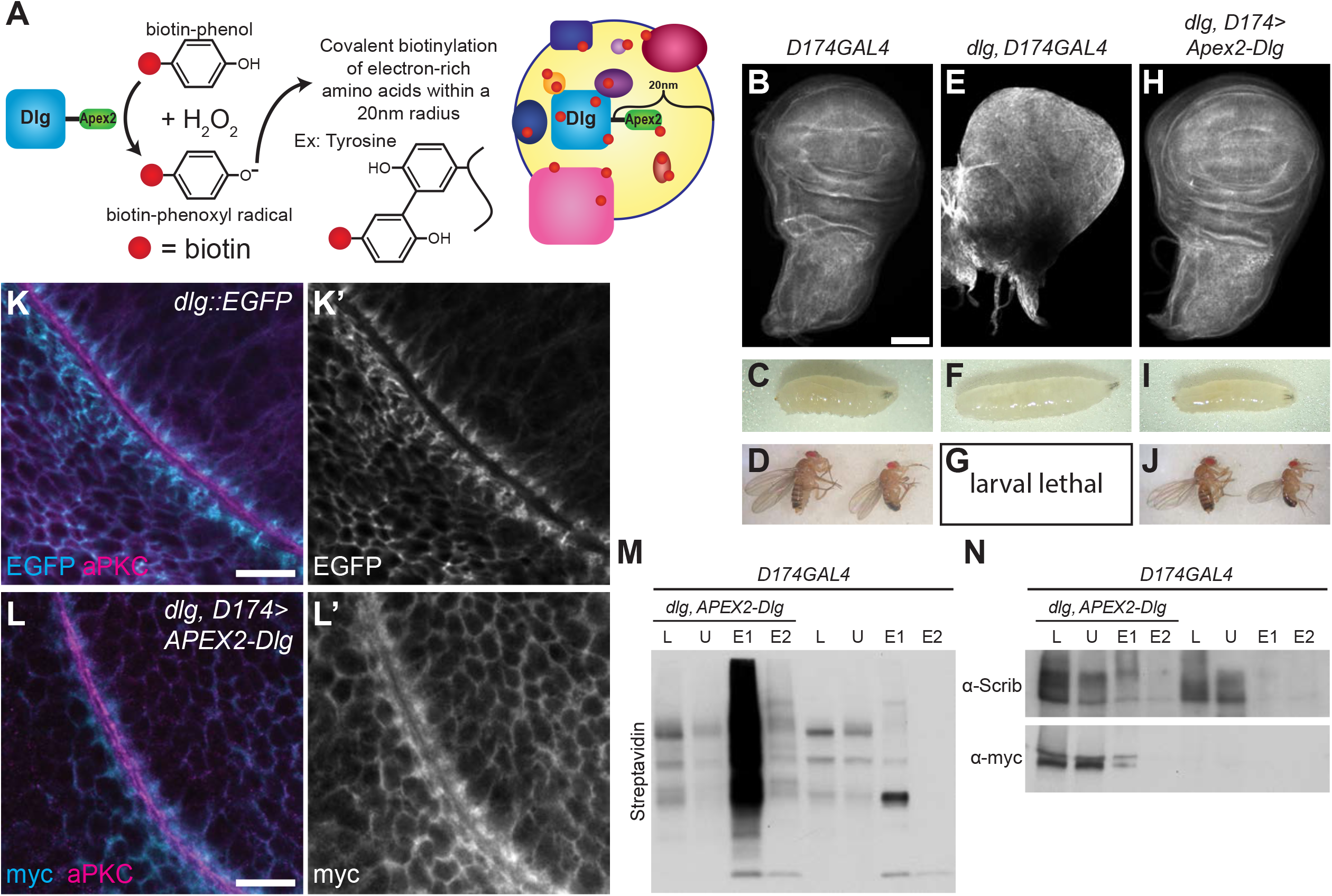
A fully functional APEX2-Dlg efficiently labels proteins with biotin. **A**. In the presence of H_2_O_2_, APEX2 catalyzes the conversion of biotin-phenol into a phenoxyl radical that can then covalently label proteins with biotin within a 20nm radius of the enzyme. **B-D:** Control *D174GAL4* 3^rd^ instar larval wing disc (**B**), larva (**C**), and adult (**D**) fly with normal size and morphology. Scale bar = 100µM. **E-G:** *dlg, D174GAL4* wing discs (**E**) form neoplastic tumors. Larvae (**F**) are overgrown and no adult flies eclose (**G**) because animals die as giant larvae. Scale bar = 100µM. **H-J:** *dlg* mutant phenotypes were rescued by *UAS-3xMyc-APEX2-Dlg* (*APEX2-Dlg*). Wing discs (**H**) have normal size and morphology, larvae are normal in size (**I**), and adult viability (**J**) and fertility are restored. Scale bar = 100µM. **K**. Endogenously tagged Dlg::EGFP reflects Dlg localization along the epithelial basolateral membrane of wing disc cells. Localization is excluded from the apical domain labeled by α-aPKC staining. Scale bar = 10µM. **L**. APEX2-Dlg (α-myc) localization recapitulated normal Dlg localization along the basolateral domain of wing disc cells and is excluded from the apical domain. Scale bar = 10µM. **M**. APEX2-Dlg efficiently labeled cellular proteins with biotin as seen by increased Streptavidin-HRP signal in experimental disc lysate (L, lane 1) compared to lysate from control discs (L, lane 5). Some biotinylated protein remained unbound to beads (U, lanes 2 and 6), but most was serially eluted off streptavidin-conjugated beads (eluate (E) 1 and 2, lanes 3, 4, 7, and 8). Biotinylation catalyzed by APEX2 was particularly apparent in these eluates. **N**. Both experimental (lane 1) and control (lane 5) lysates (L) contained Scrib, but only experimental lysate contained APEX2-Dlg (α-myc). While some Scrib and APEX2-Dlg remained unbound (and possibly unlabeled) (U, lanes 2 and 5), both were detected in the experimental but not control streptavidin-bead eluate (E1, E2, lanes 3-4 and 7-8) showing that both proteins were labeled with biotin only in the presence of the APEX2 tag.

As an entry point into understanding Scrib module function, we used an APEX2-tagged *Drosophila* Dlg to identify nearby proteins in an epithelium *in vivo*. In addition to previously proposed Dlg binding partners and other cortical proteins, there was a surprising enrichment of nuclear proteins, including the NURF complex of chromatin regulators. We demonstrate that a nuclear pool of Dlg exists in proximity to NURF members and provide evidence that NURF facilitates the growth of *dlg* tumors by activating neoplastic transcriptional programs. Our results further demonstrate the utility of proximity-based proteomics for the elucidation of the localization and function of individual proteins, particularly for multivalent scaffolding proteins.

## Results

### An APEX2-Dlg transgene for proteomics

To identify proteins enriched near Dlg in the cells of an intact epithelial sheet, we used APEX2-based *in vivo* proximity labeling. We drove an N-terminally tagged *UAS-3xMyc-APEX2-Dlg* (APEX2-Dlg) construct using a broadly but moderately expressed GAL4 driver (*D174-GAL4*) in a *dlg* null background. In these animals, APEX2-Dlg was the only Dlg protein present. APEX2-Dlg restored the morphology of *dlg* mutant discs (Figure 1B,E,H), and adult flies were rescued to viability and fertility (Figure 1C,D,F,G,I,J), demonstrating that this transgenic protein is fully functional. Because proximity-based proteomics labels not only direct binding partners but all proteins within a 20nm radius of the enzyme (Martell *et al*., 2012) (Figure 1A), it is critical that APEX2-tagged constructs localize comparably to their wild-type counterparts. Comparison to endogenously tagged Dlg::EGFP revealed that APEX2-Dlg displayed similar localization along the basolateral membranes of wing disc epithelial cells (Figure 1K-L’). We tested the enzymatic function of APEX2-Dlg by treating both control and experimental discs with hydrogen peroxide and comparing the amount of biotinylation by western blot. As expected, increased biotinylation was seen in lysate from discs expressing APEX2-Dlg as compared to control (Figure 1M), and we successfully isolated the biotinylated proteins using streptavidin beads from both control and experimental samples (Figure 1M and Supplemental Figure 1A). Finally, we assessed whether APEX2-Dlg would biotinylate proteins known to be in close proximity to endogenous Dlg. Indeed, western blotting revealed that Scrib was present in the streptavidin-bead eluate from experimental but not control samples, verifying that APEX2-Dlg was functioning as designed (Figure 1N, Supplemental Figure 1B).

### Proximity biotin labeling and mass spectrometry analysis

We next collected samples from APEX2-Dlg epithelia for mass spectrometry. Larvae were dissected to isolate the wing, haltere, and leg imaginal discs that are found together in the thorax. Samples were collected in batches, subjected to biotin labelling, and a small fraction of each post-labeling reaction lysate was reserved to verify consistent sample quality (Supp Figure 1 C,D). Batches were then pooled into three biological replicates for both the experimental and control genotype, each containing the thoracic discs of 400 larvae. Samples were tandem mass tag (TMT) labeled and then pooled for LC-MS3 (see Methods for details).

The MS results yielded a list of 485 proteins with a p-value below the statistical threshold and a log2 fold change of at least 2 between experimental and control samples (Supplemental Table 1). This list included many translation initiation and elongation factors as well as ribosomal proteins. It is possible that these proteins were labeled because APEX2-Dlg is translated from a UAS construct that is being continually produced. We therefore excluded them from further analysis, leaving a final dataset of 413 proteins (Supplemental Table 2).

We then performed cellular component Gene Ontology (GO) analysis on this dataset. Gratifyingly, enriched terms included “basolateral membrane” and “septate junction” (Figure 2A,B). The proteins that led to these terms included both Scrib and Lgl, which function with Dlg in a module to regulate polarity, as well as Cora, Nrg, FasIII, Vari, and Atpα which are junctional components whose localization is regulated by Dlg (Woods *et al*., 1996; Bilder *et al*., 2003; Oshima and Fehon, 2011; Izumi and Furuse, 2014; Lee *et al*., 2020). The dataset also included proteins previously identified as direct physical interactors of *Drosophila* Dlg (Kinesin heavy chain, Calmodulin Kinase II, 14-3-3 zeta and epsilon) (Koh *et al*., 1999; Siegrist and Doe, 2005; Nakajima *et al*., 2019). Other hits include the co-associated RNA binding proteins Caprin and Fmr1, the latter of which interacts with Lgl (Zarnescu *et al*., 2005; Baumgartner *et al*., 2013). Interestingly, “Proton-transporting V-type ATPase, V1 domain” is another enriched term in the GO analysis (Figure 2A and Supplemental Table 3), and a recent study identified an indirect physical interaction between the V-ATPase proton pump and Lgl in cultured *Drosophila* cells (Portela *et al*., 2018). Altogether, these results support the hypothesis that proximity-based proteomics can capture a snapshot of Dlg biology in living epithelia (Figure 2A and Supplemental Table 3).

**Figure 2:**
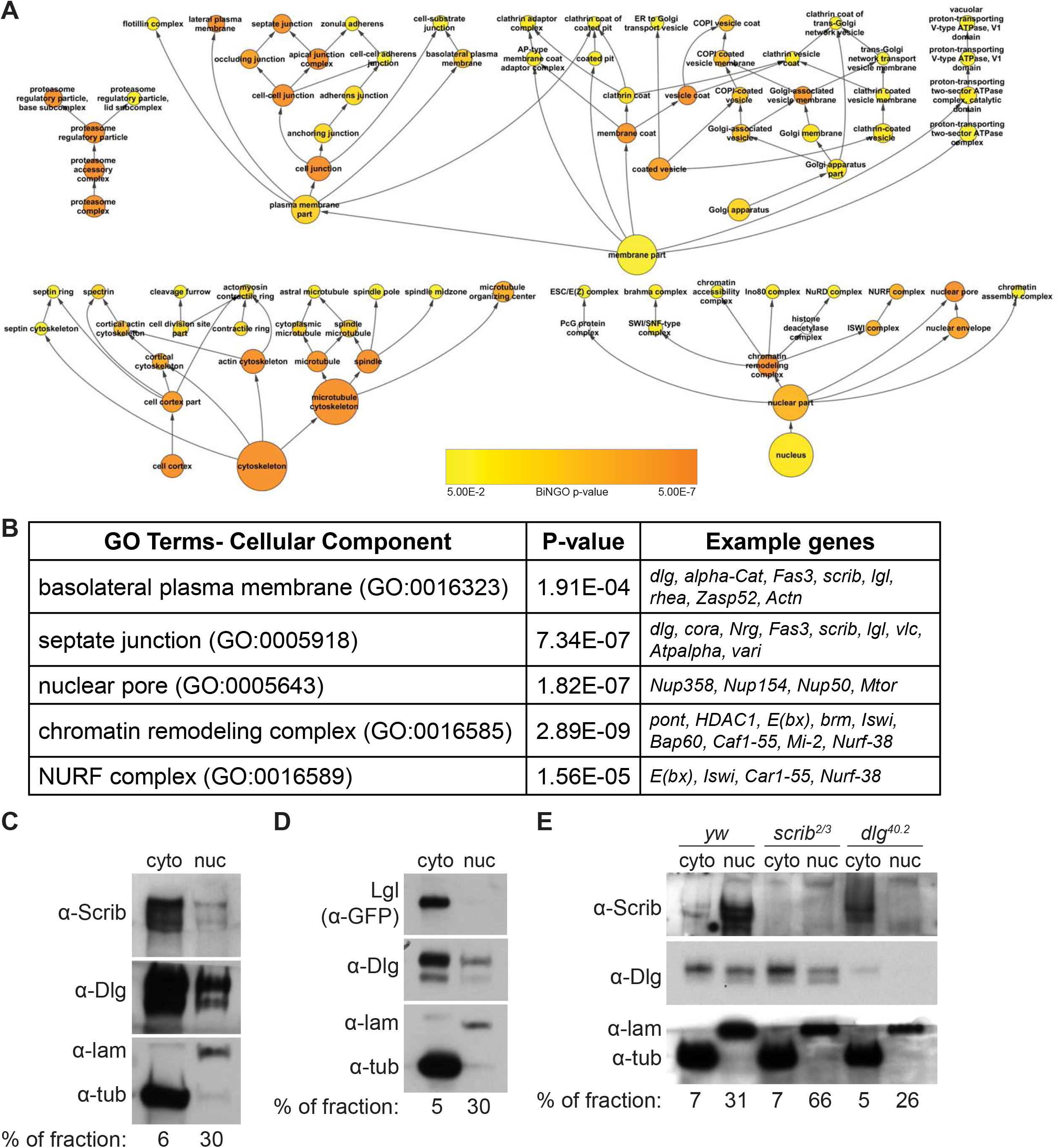
Existence of a nuclear pool of Dlg. **A**. Hierarchical map of selected enriched cellular component GO terms from APEX2-Dlg proximity biotinylation. Size of circles indicates the number of genes associated with a given term. Color of the circle indicates the p-value of enrichment for that term as indicated by the provided scale. Enriched terms include the proteasome; membrane-associated terms including regions of the cell, types of membrane associated proteins, and types of vesicles; cytoskeleton-related terms including actin and microtubule cytoskeletons; and nuclear terms. Map was pruned for viewability. Full list of enriched terms and their associated p-values is found in Supplementary Table 3. **B**. Selected enriched GO terms with their associated FDR-corrected p-values, and examples of genes from the dataset associated with each term. “Basolateral plasma membrane” and “septate junction” were expected terms, while “nuclear pore,” “chromatin remodeling complex,” and “NURF complex” were unexpected. **C-E:** Western blots of fractionated disc extracts, probed with Tubulin (α-tub) to mark the cytoplasmic fraction and Lamin (α-lam) to mark the nuclear fraction. Percentage of fraction loaded into each lane is given below; higher percent of less concentrated nuclear fractions were loaded to equalize total protein per lane. Native Scrib and Dlg are found in both cytoplasmic and nuclear fractions of disc cells (**C**), but endogenously GFP-tagged Lgl is detected only in cytoplasmic and not nuclear fraction (**D**). Dlg is still found in both the nuclear and cytoplasmic fractions of *scrib* null samples, but in *dlg* null samples, Scrib is detected only in the cytoplasmic fraction (**E**).

### Nuclear Localization of Dlg

The GO analysis also highlighted some unexpected results. Proteins annotated to the term “nucleus” were enriched in the dataset (Figure 2A,B, Supplemental Table 3). This was surprising because microscopy of fixed and immunostained tissue as well as live imaging with tagged fluorescent proteins detect Dlg localization almost exclusively at the basolateral membranes of epithelial cells (Figure 1 A,B). However, proteins associated with the GO-term “nuclear pore” were also overrepresented in the dataset and included proteins found in the pore’s cytoplasmic filaments, the central ring which spans the nuclear envelope, and the nuclear basket (Nup358, Nup155, and Nup50 and Tpr; Figure 2A,B, Supplemental Table 3). Proximity to these components is consistent with nuclear import of Dlg isoforms, all of which have molecular masses greater than 100 kDa.

We therefore investigated whether a nuclear pool of Dlg might exist in epithelia. We performed sub-cellular fractionation of wing disc lysates to generate nuclear and cytoplasmic fractions, validated by western blotting for their canonical markers Lamin and Tubulin, respectively. Strikingly, a portion of Dlg is indeed found in the nuclear fraction (Figure 2C). It is not technically feasible to quantify the exact proportion of Dlg in the nucleus because of sample loss inherent to the fractionation protocol; however, taking into account the proportion of each fraction analyzed by western blot (Figure 2C), we infer that the amount of nuclear Dlg is small compared to that found in the cytoplasm. Because Dlg, Scrib, and Lgl co-localize at the cortex and work together in many biological contexts, we asked if either Scrib or Lgl were also found in the nuclear fraction. Western blotting of fractions showed a nuclear population of Scrib (Figure 2C), but not Lgl (Figure 2D). In *dlg* null flies, this nuclear population of Scrib was lost (Figure 2E). However, in *scrib* null flies, Dlg was still found in the nuclear fraction (Figure 2E). We therefore conclude that Dlg enters the nucleus independent of Scrib and that it is required for Scrib’s nuclear localization, similar to the relationship between the proteins at the cell cortex (Bilder *et al*., 2000; Albertson and Doe, 2003; Khoury and Bilder, 2020; Ventura *et al*., 2020).

### Nuclear Dlg is in close proximity to the NURF complex

Having identified the existence of a small nuclear pool of Dlg, we considered what its function could be. In addition to “nucleus” and “nuclear pore,” the proteins associated with the term “chromatin remodeling complex” were also enriched in the APEX2-Dlg proteomic dataset (Figure 2A,B). While several chromatin remodeling complexes were over-represented, for only one were all members of the complex present in the proteomic dataset: the nucleosome remodeling factor (NURF) complex (Figure 2A,B, Supplemental Table 3).

The NURF complex is a conserved molecular machine that catalyzes, through an ATP-dependent mechanism, the sliding of nucleosomes along DNA to regulate gene expression (Badenhorst *et al*., 2002; Bouazoune and Brehm, 2006; Alkhatib and Landry, 2011; Kwon *et al*., 2016). In *Drosophila*, NURF is made up of four proteins: Iswi, Caf1-55, and Nurf-38 and E(bx) (aka NURF301) (Xiao *et al*., 2001; Alkhatib and Landry, 2011). E(bx) serves as a scaffold for the other three proteins. The NURF complex does not inherently possess sequence-specific DNA binding activity. Instead, it moves particular nucleosomes on specific target genes by binding to transcriptional regulatory proteins that themselves have DNA sequence specificity (Xiao *et al*., 2001; Alkhatib and Landry, 2011; Kwon *et al*., 2016). For example, the *Drosophila* NURF complex binds the GAGA transcription factor Trithorax-like (Trl) to move nucleosomes out of promoter regions—including those of Yki target genes—thereby facilitating transcriptional activation (Alkhatib and Landry, 2011; Oh *et al*., 2013; Kwon *et al*., 2016).

The APEX2 proteomic data suggest that Dlg is found within 20nm of the NURF complex. To verify this, we turned to a proximity ligation assay (PLA) which creates a punctate fluorescent signal when two target proteins are less than 40nm apart (Söderberg *et al*., 2006). We performed PLA using α-Dlg and α-GFP antibodies on wing discs expressing the NURF complex member E(bx) endogenously tagged with GFP. We further expressed *dlg* RNAi in the posterior compartment of the E(bx)::GFP wing discs (Figure 3A) as an internal negative control for specificity. A positive PLA signal was detected in epithelial cells from the control side of wing discs (Figure 3B) that was significantly greater than single antibody background signal (Figure 3D) and also significantly greater than signal from the *dlg-*depleted portion of the discs (Figure 3C, E). As a final control, we performed PLA on wing discs expressing Polybromo endogenously tagged with GFP. Polybromo is a member of the PBAP complex, one of two SWI/SNF chromatin remodeling complexes in *Drosophila* (Bouazoune and Brehm, 2006). We detected other SWI/SNF complex members in our MS data (Supplemental Tables 2,3) including Brm and Bap60 (Figure 2B), but not Polybromo. We therefore reasoned that Polybromo::GFP would be a stringent negative control. We did not detect PLA signal significantly above single antibody background between Dlg and Polybromo::GFP (Figure 3F).

**Figure 3:**
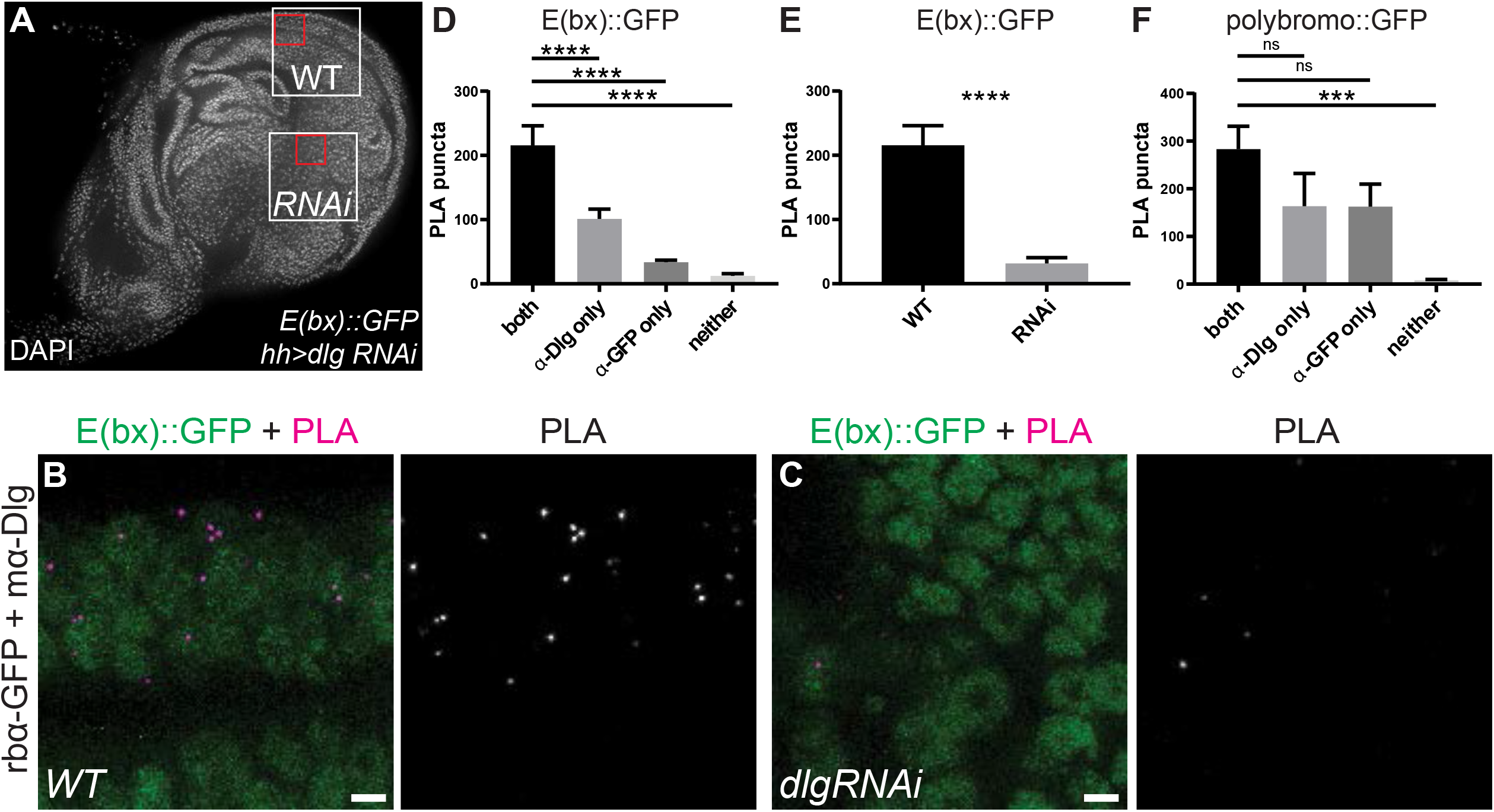
Nuclear Dlg can be detected *in vivo* in proximity to E(bx) **A**. *E(bx)::GFP* wing disc nuclei expressing *UAS-dlg-RNAi* in the posterior compartment under control of *hhGAL4*, used for PLA. Yellow boxes indicate wild-type and *dlg* knockdown areas where signal in D, E was quantified. Red boxes indicate regions shown in B,C. **B**. PLA from wild-type region; note that nearly all signal is within nuclei marked by E(bx)::GFP. Scale bar = 10µM. **C**. PLA from *dlg* RNAi region shows nearly no signal. Scale bar = 10µM. **D**. PLA signal is significantly increased between E(bx)::GFP and Dlg in WT tissue with both antibodies compared to samples with a single antibody or neither antibody. Graph displays mean with error bars of SEM. Ordinary one-way ANOVA. For ****, p<0.0001. For both, n= 14 wing discs; α-Dlg only, n= 15 wing discs; α-GFP only, n= 15 wing discs; neither, n= 12 wing discs. **E**. PLA signal is significantly increased between E(bx)::GFP and Dlg in wild-type tissue compared to *dlg* RNAi tissue. Graph displays mean with error bars of SEM. Paired, two-tailed t-test. For ****, p<0.0001. n=14 wing discs. **F**. There is no significant PLA signal between Polybromo::GFP and Dlg compared to single antibody background. Graph displays mean with error bars of SEM. Ordinary one-way ANOVA. For ns, p>0.05. For ***, p=0.0001. For both, n= 16 wing discs; α-Dlg only, n= 13 wing discs; α-GFP only, n= 13 wing discs; neither, n= 16 wing discs.

An advantage of PLA in this context is its sensitivity, which allows small numbers of protein molecules to be visualized via microscopy with good spatial resolution. The PLA signal detected between Dlg and E(bx)::GFP appeared in the nuclei of epithelial cells (Figure 3B). Because Dlg has a well-established role in regulating spindle orientation during cell division (Albertson and Doe, 2003; Johnston *et al*., 2009; Bergstralh *et al*., 2013; Nakajima *et al*., 2019), we considered that the nuclear Dlg signal from both MS and cell fractionation could derive exclusively from dividing cells. However, the PLA signal was even across cells and not limited to those undergoing mitosis. Thus, in addition to confirming a population of Dlg near the NURF complex, this method also enabled the detection of endogenous Dlg in the nuclei of intact cells, for the first time to our knowledge. These data corroborate the evidence from MS and sub-cellular fractionation that a nuclear pool of Dlg exists and lies specifically near the NURF complex.

### The NURF complex is required for overgrowth of *dlg* tumors

We next sought to determine if there was a functional connection between Dlg and the NURF complex. Depleting *dlg* with RNAi from the boundary between anterior and posterior compartments of the wing disc using a conditionally active *ptc*^*ts*^*GAL4* (see Methods) can cause neoplastic overgrowth in the hinge regions both proximal and distal to the pouch (Figure 4D). Depleting the NURF complex components *E(bx)* or *iswi* in otherwise WT discs using the same GAL4 driver has only minor effects, inducing limited apoptosis in the pouch that leads to a slight narrowing of the stripe (Figure 4A-C). Strikingly, co-expressing RNAi against either NURF complex component with *dlg* RNAi rescued the overgrowth of *dlg*-depleted cells (Figure 4E-F). Reduction in *dlg* RNAi tumor size by RNAi against NURF components was not specific to the *ptc*^*ts*^*GAL4* driver, as similar results were seen for tumors in the posterior compartment of wing discs induced using *hhGal4* (Supplemental Figure 2D-F). Again in this system, expressing either NURF RNAi alone had little to no effect on tissue morphology or size although these animals showed a small developmental delay (Supplemental Figure 2A-C). Although RNAi against NURF complex components partially rescued *dlg* RNAi tumor overgrowth, we did not observe rescue of cell polarity or tissue architecture (Supplemental Figure S3A-D).

**Figure 4:**
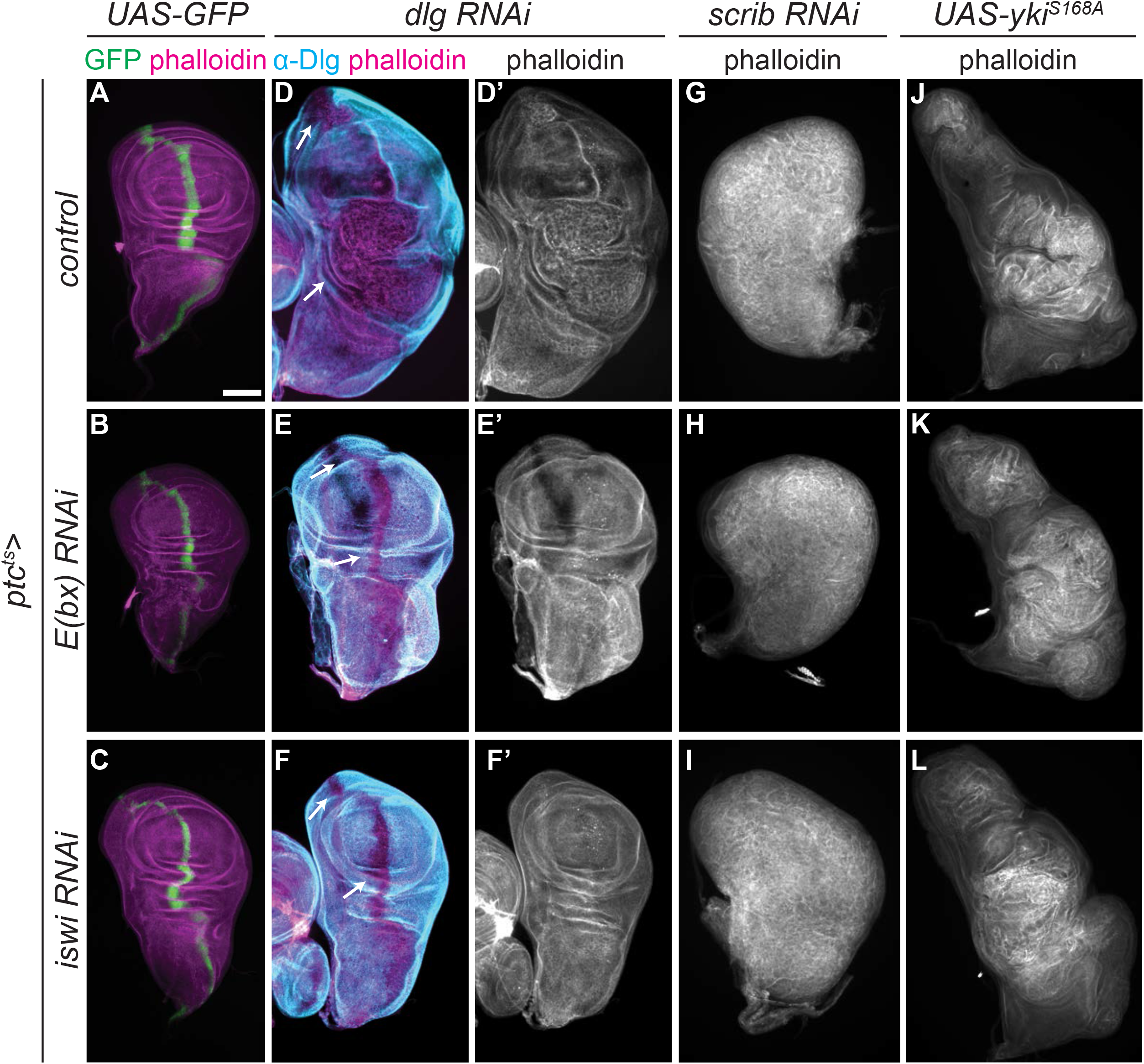
Overgrowth of *dlg*-depleted epithelia requires NURF complex. **A-C**. *ptc*^*ts*^*GAL4* drives expression of GFP along the AP compartment boundary (**A**). Discs expressing RNAi against NURF complex members *E(bx)* (**B**) or *iswi* (**C**) in the Ptc stripe have normal size and morphology. **D-F**. Depleting *dlg* in the Ptc stripe caused neoplastic overgrowth in both proximal and distal hinge regions (**D**, arrows). Overgrowth was rescued by either *E(bx)* RNAi (**E**) or *iswi* RNAi (**F**). **G-I**. Depleting *scrib* with RNAi in the Ptc stripe also caused neoplastic tumors (**G**), but tumor formation and growth were unaffected by co-expression of either *E(bx)* (**H**) or *iswi* (**I**) RNAi. **J-L**. Expressing the constitutively active YkiS168A mutant in the Ptc stripe caused hyperplastic overgrowth of wing discs (**J**). Yki-driven overgrowth was unaffected by co-expression of either *E(bx)* (**K**) or *iswi* (**L**) RNAi. All scale bars = 100µM.

We tested the specificity of the requirement of NURF complex for tumor growth. A constitutively-active form of Yki (S168A) induces strong hyperplastic overgrowth when expressed in the *ptc*^*ts*^*GAL4* stripe (Figure 4J) but this growth was not rescued by *E(bx)* or *iswi* RNAi (Figure 4K,L). Even though there is also a nuclear pool of Scrib (Figure 2C,E), overgrowth in *scrib* RNAi tumors was also not rescued by RNAi against either NURF component (Figure 4G-I). Thus, although the reduction of tumor size by NURF RNAi is not a general characteristic of all tumors.

### NURF complex promotes Yki target-gene expression in *dlg* tumors

Transcriptional changes that drive neoplastic tumor growth in *Drosophila* are driven by a signaling network involving JNK-mediated regulation of the Fos transcription factor and aPKC-mediated regulation of Yki (Kulshammer *et al*., 2015; Atkins *et al*., 2016). Because the NURF complex, via its interaction with Trl, participates in activation of Yki target genes (Oh *et al*., 2013), we asked if this might account for NURF complex role in promoting *dlg* tumor growth. NURF RNAi-mediated suppression of *dlg*-depleted tumor growth did not rescue epithelial architecture (Supplemental Figure 2), but it caused significant suppression of a reporter for Yki activity, *ex-LacZ* (McCartney *et al*., 2000; Hamaratoglu *et al*., 2006). While NURF RNAi alone had no effect on expression of *ex-LacZ* in otherwise WT tissue (Figure 5A-C), *ex-LacZ* expression was reduced to normal levels in *dlg* RNAi tissue co-expressing either NURF RNAi (Figure 5D-F). However, in *scrib* RNAi tumors, where overgrowth is not suppressed, *ex-LacZ* levels are unchanged by NURF RNAi (Figure 5G-I). Thus, the ability of NURF RNAi to limit tumor overgrowth correlates with the extent to which it limits Yki target gene activation in that tumor type. The failure of NURF RNAi to reduce the overgrowth caused by Yki^S168A^ (Figure 4) suggests that the NURF complex is not absolutely required for Yki to drive proliferation, and that strong, constitutive activation of Yki can overcome an unfavorable chromatin environment to drive gene expression. Altogether, these data support a model where nuclear Dlg negatively regulates the NURF complex to limit Yki driven growth.

**Figure 5:**
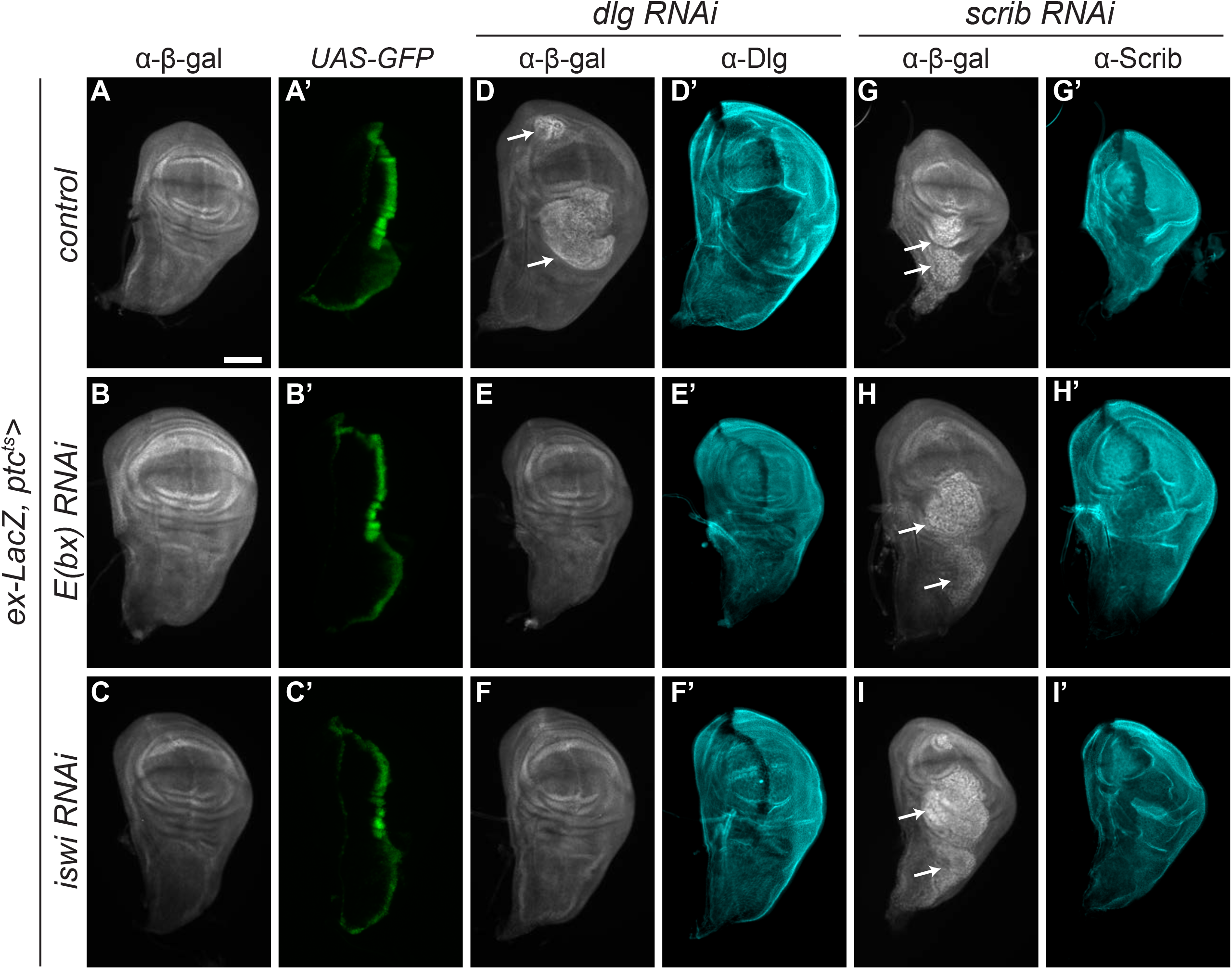
NURF RNAi rescues Yki target gene expression in *dlg* tumors. **A-C**. *ex-LacZ* expression (α-β-gal) in control wing discs (**A**) was unaffected by either *E(bx)* (**B**) or *iswi* (**C**) RNAi. **D-F**. *ex-LacZ* levels were elevated in neoplastically overgrown regions of *dlg* RNAi discs. (**D**, arrows). *E(bx)* (**E**) or *iswi* (**F**) RNAi coexpression rescued both overgrowth and *ex-LacZ* levels. **G-I**. *ex-LacZ* levels were also elevated in neoplastically overgrown areas of *scrib* RNAi discs (**G**). Neither overgrowth nor *ex-LacZ* levels were rescued with co-expression of *E(bx)* (**H**) or *iswi* (**I**) RNAi. The phenotype here was made less severe than in Figure 4G-I by reducing *ptc*^*ts*^*GAL4* activity time, to facilitate comparison of normal to tumorous tissue. All scale bars = 100µM.

### Nuclear localization of Dlg involves sequences outside of consensus NLSs

In order to explore a function for nuclear Dlg, we investigated potential nuclear localization signals (NLSs). Prediction algorithms consistently identified two regions enriched in basic amino acids that are characteristic of recognized NLSs (Figure 6A). The first lies at the C-terminus of PDZ1, while the second is in the so-called E-F region at the N-terminus of the HOOK domain, which itself lies between the SH3 and GUK domains. Both are conserved in the human homolog hDlg1, and for the latter, experimental evidence consistent with an NLS has been demonstrated. Transgenic constructs have previously deleted either the E-F or the entire HOOK domain; each results in mutant Dlg proteins that show strongly increased nuclear localization visible by immunohistochemistry (Hough *et al*., 1997; Lu *et al*., 2021). Since these data indicate that the E-F region cannot constitute the sole NLS of Dlg, we mutated basic residues within the predicted NLSs in both the E-F region as well as PDZ1 to alanine (Dlg^2XNLS>A^, Figure 6A).

**Figure 6:**
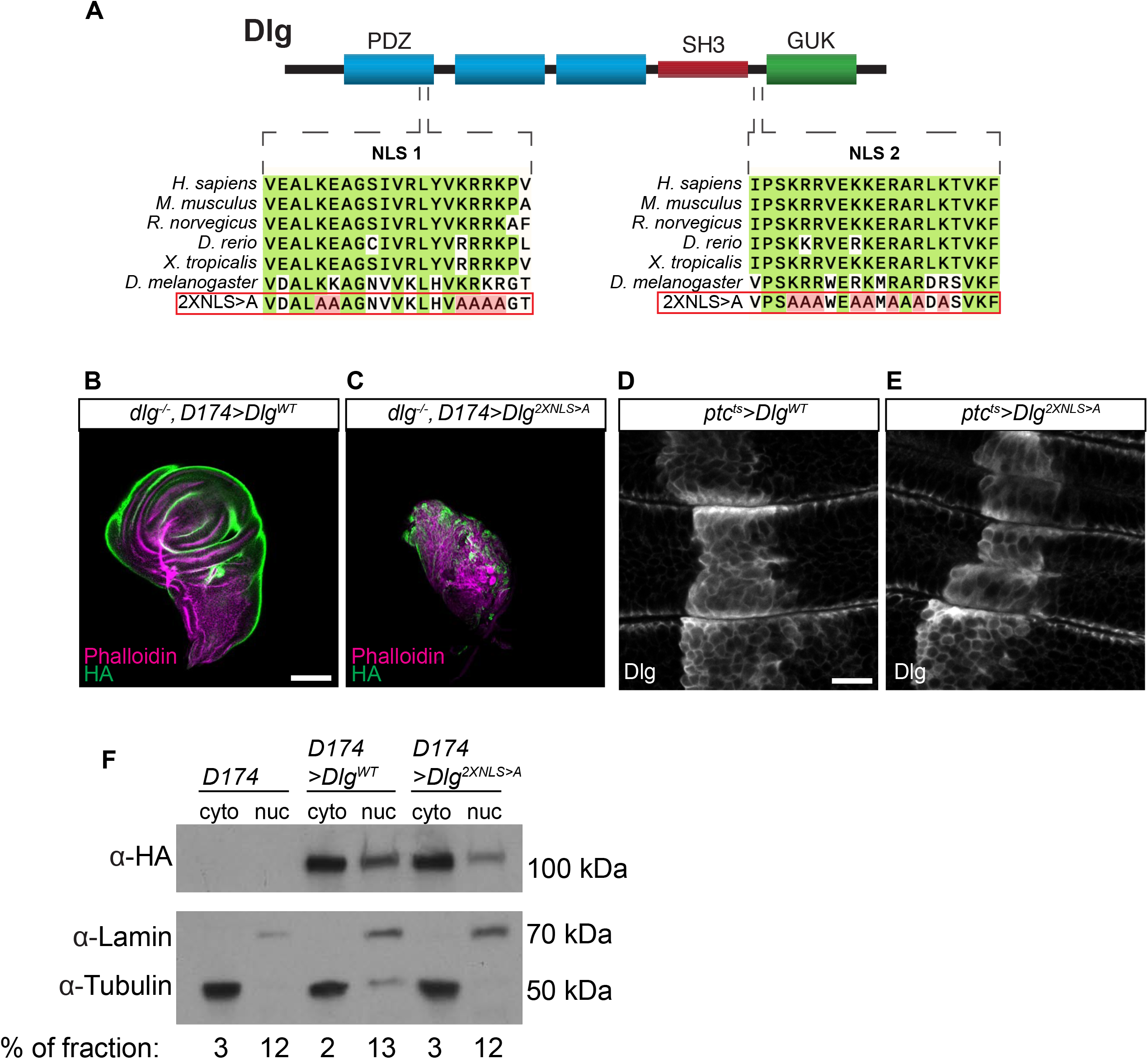
Dlg nuclear localization involves sequences outside of predicted NLSs. **A**. Diagram of Dlg protein, with predicted nuclear localization signals (NLSs) shown in relation to conserved domains. Conservation of NLSs is shown in alignment of Dlg homologs. Mutations in Dlg^2XNLS>A^ are shown in red. **B, C**. *dlg* null mutant wing discs are rescued by expression of Dlg^WT^ (**B**) but not Dlg^2XNLS>A^ (**C**) **D, E**. Localization of Dlg^WT^ (**D**) under *ptc-GAL4* is comparable to Dlg^2XNLS>A^ (**E**), although the latter shows more cytoplasmic staining. Transgenes are detected by anti-Dlg staining: compare to endogenous protein localization neighboring the stripe. **F**. Western blots of fractionated disc extracts expressing transgenic Dlg proteins, detected by α-HA. Blots are also probed with Tubulin (α-tub) to mark the cytoplasmic fraction and Lamin (α-lam) to mark the nuclear fraction. Dlg^WT^ and Dlg^2XNLS>A^ are both found in the nuclear as well as the cytoplasmic fraction.

When overexpressed in *dlg* mutant discs or follicle cells, Dlg^2XNLS>A^ had no detectable rescuing activity compared to expression of WT Dlg (Figure 6B,C, Supplemental Figure 4H-J). Both Dlg^2XNLS>A^ and matched WT Dlg transgenic proteins exhibited membrane and cytoplasmic localization in disc and follicle cells (Figure 6E, Supplemental Figure 4A-D). Quantitation of membrane localization in follicle cells showed that, while Dlg^2XNLS>A^ was enriched at the membrane (PM Index >1), its localization was reduced compared to WT Dlg (Supplemental Figure 4A-D,G). We considered whether this might be due to disruption of electrostatic membrane binding, since the mutated residues in NLS2 of Dlg^2XNLS>A^ partially overlap with a recently described membrane binding polybasic region (Figure 6A)(Lu *et al*., 2021). However, localization was further reduced in *scrib-*depleted cells (Supplemental Figure 4E,F), while Dlg constructs that lack electrostatic membrane binding are insensitive to *scrib* depletion (Lu *et al*., 2021). Most importantly, to determine if mutation of both predicted NLS altered Dlg’s ability to enter the nucleus, we carried out subcellular fractionation studies on protein extracted from imaginal discs. Dlg^2XNLS>A^ protein could still be detected in the nuclear fraction, similarly to the matched overexpressed WT Dlg (Figure 6F). We conclude that nuclear entry of Dlg can involve sequences outside of the two predicted NLSs.

## Discussion

Here we report the use of an *in vivo* proximity-based biotin labeling approach to investigate new biological functions of the polarity-regulating tumor suppressor Dlg. Our strategy used transgenic replacement of Dlg by an APEX2-tagged version, coupled with mass spectrometry of subsequently tagged proteins; critically, we carried this out in native epithelial tissue. The resultant MS data led to the discovery of a nuclear pool of native Dlg that is not apparent by microscopy but that nevertheless lies near the NURF chromatin remodeling complex. We further found that NURF is required for the overgrowth of epithelia lacking Dlg and suggest that this may be due to the role of NURF in activating pro-proliferative *yki* target genes.

To our knowledge, our data provide the first demonstration of endogenous *Drosophila* Dlg in the nucleus. Dlg is a member of the membrane-associated guanylate kinase (MAGUK) family of proteins that are generally found at cell-cell junctions, but nuclear localization of many MAGUKs has also been observed including hDlg1 (Mantovani and Banks, 2003; Roberts *et al*., 2007; Narayan *et al*., 2009) as well as ZO-1 (Gottardi *et al*., 1996; González-Mariscal *et al*., 1999), ZO-2 (Islas *et al*., 2002), CASK1 (Hsueh *et al*., 2000), MAGI-2 (Dobrosotskaya *et al*., 1997), and Nagie Oko (Bit-Avragim *et al*., 2008). Due to their size, movement of MAGUKs into and out of the nucleus must involve active transport via NLS and nuclear export signals (NES) respectively, consistent with the presence of several importin proteins as well as the nuclear import regulator Ran in our Dlg proximity biotinylation dataset (Supplemental Tables 1,2). The predicted NLS in the Dlg E-F/HOOK region is conserved in hDlg1. The SH3-HOOK-GUK region undergoes an intramolecular interaction that regulates exposure of sequences that can mediate several Dlg functions (Nix *et al*., 2000; McGee *et al*., 2001; Qian and Prehoda, 2006; Marcette *et al*., 2009; Newman and Prehoda, 2009; Zeng *et al*., 2017; Rademacher *et al*., 2019; Lu *et al*., 2021); the increased nuclear localization of transgenic Dlg or hDlg1 deleted for the GUK domain raises the possibility that nuclear entry could be one of these (Thomas *et al*., 2000; Kohu *et al*., 2002; Lu *et al*., 2021). The predicted NLS in Dlg PDZ1 is conserved not only in hDlg1 but also in the MAGUKs ZO-1, ZO-2 and their *Drosophila* homolog Polychaetoid. We were unable to predict an NES, although such sequences have been detected in vertebrate ZO-1 (Islas *et al*., 2002), ZO-2 (Islas *et al*., 2002; Jaramillo *et al*., 2004; González-Mariscal *et al*., 2006), and Nagie Oko (Bit-Avragim *et al*., 2008). However, various transgenically-expressed truncations of Dlg localize strongly to nuclei (Hough *et al*., 1997; Thomas *et al*., 2000; Bachmann *et al*., 2004; Lu *et al*., 2021), including one that deletes the E-F region itself. Additionally, coding exons that are alternatively spliced in a neural-specific isoform of Dlg are sufficient to drive nuclear localization; these protein sequences are conserved with hDlg1 (Bachmann *et al*., 2004). Clearly, MAGUKs have evolved and maintained multiple mechanisms of nuclear entry and exit; our data demonstrate that this may be true even for proteins where a nuclear pool is not visible by microscopy.

We attempted to test the nuclear function of *Drosophila* Dlg by mutating two conserved and predicted NLSs; however, this failed to abolish nuclear entry. This result agrees with a recently published Dlg transgene that deletes the three PDZ domains while also mutating 15 basic residues in and adjacent to the predicted E-F region NLS to glutamine; nuclear localization of the mutant protein is visible, again emphasizing that an additional NLS must reside outside of the consensus sequences (Lu *et al*., 2021). Interestingly, Dlg^2XNLS>A^ provided no rescuing activity, consistent with the deleterious effects of mutations in the HOOK domain (Hough *et al*., 1997; Lu *et al*., 2021). Previous work shows that PDZ and GUK domains are not absolutely required for Dlg function, while the SH3 and HOOK domains are (Hough *et al*., 1997; Thomas *et al*., 2000; Khoury and Bilder, 2020; Lu *et al*., 2021). Our results hint at a critical role for the 8 basic amino acids in the E-F region within HOOK that is independent of electrostatic membrane binding; these can be the subject of future experiments.

Nakajima and Gibson recently used coimmunoprecipitation to isolate Dlg-associated proteins from embryos. Their dataset identified several proteins involved in nuclear pore traffic (Ran, CRM1, NUP358) but not NURF components (Nakajima *et al*., 2019). Similarly, we were unable to co-immunoprecipitate (Co-IP) Dlg and NURF complex member E(bx) (data not shown). It is possible that Dlg and E(bx) associate through an interaction that is not stable enough to pull down or that Dlg may interact with another member of the NURF complex (for which reagents for co-IP were not available) or through an intermediary protein. Any of these possibilities would highlight the utility of APEX2 fusion proteins to facilitate the *in vivo* detection of weak and/or transient interactions that are hard to discover by other methods. In the case of the Scrib module, they could provide a path forward to identifying relevant interactors, which has been a major obstacle in understanding the biology of these key polarity-regulating tumor suppressors.

It has long been observed that Scrib, Dlg, and Lgl are required to maintain the proper transcriptional state of epithelial tissues (Hariharan and Bilder, 2006; Grzeschik *et al*., 2010; Zhu *et al*., 2010; Doggett *et al*., 2011; Sun and Irvine, 2011; Bunker *et al*., 2015). Our observations that Dlg is near the NURF complex in the nucleus and that the NURF complex is required for Yki-driven overgrowth of *dlg* tumors suggest that Dlg may regulate transcription through a much more direct mechanism than previously thought. Several other MAGUKs physically interact with transcription factors, including ZO-2 with Jun, Fos, C/EBP (Betanzos *et al*., 2004), Myc (Huerta *et al*., 2007), and YAP (Oka *et al*., 2010) (the vertebrate homolog of Yki), while CASK1 can form a trimeric complex in the nucleus with the T-box transcription factor Tbr-1 and CINAP, a nucleosome assembly protein that facilitates chromatin remodeling (Hsueh *et al*., 2000; Wang *et al*., 2004). Thus, the association between nuclear *Drosophila* Dlg and the NURF complex extends data about other nuclear MAGUK proteins. Still, as we were unable to separately test the functions of cytoplasmic and nuclear Dlg pools, we cannot definitively conclude that the latter drives the observed growth phenotypes. We also do not know how Dlg proteins move through the pore; whether the nuclear population is a small, stable pool or a transient, dynamic one; or the signals or cellular states that cause their nuclear localization. Nonetheless, our data indicate that a comprehensive understanding of this critical protein may have to include its function not just at the plasma membrane but also in the nucleus.

## Methods

### Drosophila stocks and Genetics

*Drosophila melanogaster* stocks were raised on cornmeal molasses food. Experimental crosses were raised at 25°C unless otherwise noted. Fly lines used are listed in Supplemental Table 4. For experiments using *ptc*^*ts*^*GAL4*, crosses were started at 25°C. Adults were removed after 48 hours then after another 24 hours, vials were shifted to 29°C to induce Gal4 expression. Wing discs were dissected from wandering third instar larvae starting 72 hours after temperature shift and continuing every 24 hours thereafter for 1-3 additional days to track tumor development. For clonal GAL4 expression in follicle cells, larvae were heat shocked for 13 minutes at 37°C 120 hours after egg deposition (AED). For follicle cell MARCM experiments, larvae were heat shocked for 1 hour on 3 consecutive days starting at 120 hours AED. Ovaries were dissected from adult females fed on yeast for 3 days after eclosion. The sequences for *UAS-3xMyc-APEX2-Dlg, UAS-Dlg-3xHA* and *UAS-Dlg*^*2XNLS>A*^*-3xHA* are presented in Supplemental Table 5. *UAS-3xMyc-APEX2-Dlg* contains the *dlgA* coding sequence with the *dlg S97*-specific exon using a cDNA provided by Ulrich Thomas; it was cloned into a Gateway N-terminal *3xMyc-APEX2* destination vector. *UAS-Dlg-3xHA* was cloned into the *pUAST-attB* vector and contains a C-terminal flexible linker followed by a 3xHA tag. *UAS-Dlg*^*2XNLS>A*^*-3xHA* was cloned from *UAS-Dlg-3xHA* using Gibson assembly (NEB). All transgenes were inserted into the *attP40* site. Primer sequences are given in Supplemental Table 5.

### Immunofluorescence and Microscopy

Wandering third-instar larval imaginal discs were dissected in PBS and fixed for 20 minutes in 4% PFA. Samples were rinsed in PBS. Follicles were dissected in Schneider’s medium containing 15% FBS and fixed for 20 minutes in 4% PFA. Primary and secondary antibodies were diluted in PBST (0.1% Triton X-100) with 4% NGS (Gibco) and 1% BSA (Gibco). Primary antibodies (Supplemental Table 4) were incubated with samples overnight at 4°C. Secondary fluorophore-conjugated antibodies (Molecular Probes) were diluted 1:400 and incubated for 2 hours at room temperature. Phalloidin and DAPI incubated with samples for 20 minutes in PBS. Images were captured on a Zeiss LSM700 scanning confocal microscope or a Zeiss Axio Imager M2 with Apotome 2 with Plan Apochromat 20x/NA 0.8, LD C-Apochromat 40x/NA 1.1 W and Plan Apochromat 63x/NA 1.4 oil objectives at 1024×1024 pixels with 2 line averages.

Dlg cortical enrichment was quantified as described in (Lu *et al*., 2021). For single cells in en face confocal sections, the fluorescent signal intensity of at the membrane and in the cytoplasm were quantified in Fiji (Schindelin *et al*., 2012) using rectangular ROIs of fixed 1.17µm width, approximately the thickness of the cell cortex. The ratio of membrane:cytoplasmic intensity was calculated to give the “plasma membrane index” (PM index). Average PM indices were calculated for all cells per genotype.

### Western Blots

Protein concentrations in samples were measured by BCA protein assay (Pierce). Proteins were electrophoresed at 150V for one hour through 7.5% or 4-20% mini-PROTEAN TGX gels (Bio-Rad) and blotted at 300 mA for one hour onto PVDF membranes. Membranes were blocked in 3% BSA in TBS-T for one hour. All antibodies were incubated with membrane in blocking solution. Primary antibodies were incubated overnight at 4°C. Streptavidin-HRP and HRP-conjugated secondary antibodies were incubated with the membrane for two hours at room temperature. Blots were developed with standard ECL reagents (Advansta).

### Sample preparation and biotin labeling for Mass Spectrometry

Larvae were reared at room temperature (21-23°C). Thoracic discs were dissected in chilled labeling media: Schneider’s medium (Gibco) containing 1% Pen/Strep (Caisson Labs), 10% FBS (Gibco), 500µM biotin-phenol (aka biotinyl tyramide, AdipoGen Life Sciences), and 2mM probenecid (Thermo). Samples were incubated at room temperature with nutation for 30 minutes. Labeling media was removed. Samples were incubated in 1mM hydrogen peroxide for 1 minute. Samples were then washed three times in quenching buffer [5mM Trolox (Sigma), 10mM sodium azide (Sigma), and 10mM sodium ascorbate (Sigma) in PBS] and three times in PBS. Samples were lysed in RIPA buffer [50mM Tris-HCl, pH8.0, 150mM NaCl, 1% NP-40, 0.5% sodium deoxycholate, 0.1% SDS and protease inhibitors (Pierce mini-tablets)] using a pellet pestle motor and a polypropylene pestle. All solutions were pre-chilled on ice. Lysates were spun at 14,000 rpm in a table-top microfuge for 10 minutes at 4°C to remove debris, and supernatants were saved as final sample lysate. A 5µL lysate sample was reserved for western blot analysis. Remainder was incubated with streptavidin-conjugated magnetic beads (Pierce) for one hour at room temperature. Beads were washed twice in TBS with 0.1% Tween20, then three times with RIPA buffer. For western blot analysis of pull-down of biotinylated proteins, beads were boiled for 5 minutes in 60µL SDS-PAGE sample buffer: NuPAGE LDS buffer (Thermo) with 20mM DTT (Sigma) and 2mM biotin (Thermo). Supernatant was saved as eluate 1 (E1). Beads were then boiled for 5 minutes in 40µL sample buffer and supernatant was saved as eluate 2 (E2). For mass spectrometry, labeled lysates were prepared in batches and stored at -80°C until all samples had been collected. Samples were then thawed, pooled into three replicates from 400 larvae each, and biotinylated proteins were isolated as described above. Protein-bound beads were kept in PBS at 4°C. Mass spectrometry, including remaining sample prep, was performed by the UC Davis Mass Spectrometry Facilities.

Proteins on beads were received and the buffer was exchanged with 4 washes of 50mM TEAB (Tri Ethyl Ammonium Bicarbonate). The proteins were then digested off the beads overnight with trypsin at room temperature. The following day, the supernatant was removed, and the beads were washed with 50mM TEAB and pooled with the supernatant. The peptides in all six sample were quantified using Pierce Fluorescent Peptide assay (Thermo Scientific).

### TMT Labeling

Based on the Fluorescent Peptide assay, the volume for 20 μg of the most concentrated sample was determined, and equal volumes of each sample were diluted with 50mM TEAB to 25 μl per replicate. Each sample was labeled with TMT 6 Plex Mass Tag Labeling Kit (Thermo Scientific). Briefly, 20 μl of each TMT label (126-131) was added to each digested peptide sample and incubated for an hour. The reaction was quenched with 1μl of 5% Hydroxylamine and incubated for 15 minutes. All labeled samples were then mixed together and lyophilized to almost dryness. The TMT labeled sample was reconstituted in 2% Acetonitrile 0 .1% TFA and desalted with a zip tip.

### LC-MS3

LC separation was done on a Dionex nano Ultimate 3000 (Thermo Scientific) with a Thermo Easy-Spray source. The digested peptides were reconstituted in 2% acetonitrile /0.1% trifluoroacetic acid, and 5µl of each sample was loaded onto a PepMap 100Å 3U 75 um x 20 mm reverse phase trap where they were desalted online before being separated on a 100 Å 2U 50 micron x 150 mm PepMap EasySpray reverse phase column. Peptides were eluted using a 90-minute gradient of 0.1% formic acid (A) and 80% acetonitrile (B) with a flow rate of 200nL/min. The separation gradient was run with 2% to 5% B over 1 minute, 5% to 10% B over 9 minutes, 10% to 20% B over 27 minutes, 20% to 35% B over 10 minutes, 35% to 99%B over 10 minutes, a 2 minute hold at 99%B, and finally 99% to 2%B held at 2% B for 5 minutes.

### MS3 Synchronous Precursor Selection Workflow

Mass spectra were collected on a Fusion Lumos mass spectrometer (Thermo Fisher Scientific) in a data-dependent MS3 synchronous precursor selection (SPS) method. MS1 spectra were acquired in the Orbitrap, 120K resolution, 50ms max inject time, 5 x 105 max inject time. MS2 spectra were acquired in the linear ion trap with a 0.7Da isolation window, CID fragmentation energy of 35%, turbo scan speed, 50 ms max inject time, 1 x 104 AGC and maximum parallelizable time turned on. MS2 ions were isolated in the ion trap and fragmented with an HCD energy of 65%. MS3 spectra were acquired in the orbitrap with a resolution of 50K and a scan range of 100-500 Da, 105 ms max inject time and 1 x 105 AGC.

### MS3 SPS Workflow

Database searching: Tandem mass spectra were extracted by Proteome Discoverer 2.2. Charge state deconvolution and deisotoping were not performed. All MS/MS samples were analyzed using SequestHT (XCorr Only) (Thermo Fisher Scientific, San Jose, CA, USA) in Proteome Discoverer 2.2.0.388). Sequest (XCorr Only) was set up to search uniprot-proteome-3AUP000000803.fasta (unknown version, 21134 entries) and an equal number of decoy sequences, assuming the digestion enzyme trypsin. Sequest (XCorr Only) was searched with a fragment ion mass tolerance of 0.60 Da and a parent ion tolerance of 10.0 PPM. Carbamidomethyl of cysteine and TMT6plex of lysine were specified in Sequest (XCorr Only) as fixed modifications. Deamidated of asparagine, oxidation of methionine and acetyl of the n-terminus were specified in Sequest (XCorr Only) as variable modifications.

Criteria for protein identification: Scaffold (version Scaffold_4.8.4, Proteome Software Inc., Portland, OR) was used to validate MS/MS based peptide and protein identifications. Peptide identifications were accepted if they could be established at greater than 74.0% probability to achieve an FDR less than 1.0% by the Scaffold Local FDR algorithm. Protein identifications were accepted if they could be established at greater than 35.0% probability to achieve an FDR less than 1.0% and contained at least 2 identified peptides. This resulted in a peptide decoy FDR of 0.31% and a Protein Decoy FDR of 0.8%. Protein probabilities were assigned by the Protein Prophet algorithm (Nesvizhskii, Al et al Anal. Chem. 2003;75(17):4646-58). Proteins that contained similar peptides and could not be differentiated based on MS/MS analysis alone were grouped to satisfy the principles of parsimony. Proteins sharing significant peptide evidence were grouped into clusters.

Quantitative data analysis: Scaffold Q+ (version Scaffold_4.8.4, Proteome Software Inc., Portland, OR) was used to quantitate Label Based Quantitation (TMT) peptide and protein identifications. Peptide identifications were accepted if they could be established at greater than 74.0% probability to achieve an FDR less than 1.0% by the Scaffold Local FDR algorithm. Protein identifications were accepted if they could be established at greater than 35.0% probability to achieve an FDR less than 1.0% and contained at least 2 identified peptides. Protein probabilities were assigned by the Protein Prophet algorithm (Nesvizhskii, Al et al Anal. Chem. 2003;75(17):4646-58). Proteins that contained similar peptides and could not be differentiated based on MS/MS analysis alone were grouped to satisfy the principles of parsimony. Proteins sharing significant peptide evidence were grouped into clusters. Channels were corrected by correction factors supplied by the manufacturer in all samples according to the algorithm described in i-Tracker (Shadforth, I et al BMC Genomics 2005;6 145-151). Normalization was performed iteratively (across spectra) on intensities, as described in Statistical Analysis of Relative Labeled Mass Spectrometry Data from Complex Samples Using ANOVA (Oberg, Ann L. et al., Journal of proteome research 7.1 (2008): 225–233). Medians were used for averaging. Spectra data were log-transformed, pruned of those matched to multiple proteins, and weighted by an adaptive intensity weighting algorithm. Of 3635 spectra in the experiment at the given thresholds, 3247 (89%) were included in quantitation. Differentially expressed proteins were determined by applying Permutation Test with unadjusted significance level p < 0.05 corrected by Benjamini-Hochberg.

### Data Availability

Raw data, mzML and Scaffold results are available from the MassIVE proteomics repository (MSV000087186) and Proteome Exchange (PXD025378).

### GO analysis

Cellular component Gene Ontology analysis was performed using the BiNGO plug-in for Cytoscape using all genes in the *Drosophila melanogaster* genome as a reference set. User selected settings were as follows: hypergeometric statistical test; Benjamini-Hochberg false discovery rate (FDR) used to correct P-values; significance level set to 0.05.

### Multiple Sequence Alignment

Multiple sequence alignment of Dlg NLS sequences was made with Clustal Omega (Madeira *et al*., 2019). Aligned sequences were visualized using SnapGene Viewer. The following protein sequences were used: *H. sapiens*: UniProt Q12959, *M. musculus*: UniProt Q811D0, *R. norvegicus*: UniProt Q62696, *D. rerio*: UniProt E7FAT1, *X. tropicalis*: UniProt Q28C55, *D. melanogaster*: UniProt P31007.

### Subcellular Fractionation

60-70 wandering third instar larvae were dissected in PBS. Cells were lysed using 5 strokes with the small pestle in a Dounce homogenizer in: 10mM HEPES, pH7.6, 10mM KCl, 1.5mM MgCl_2_, .5 mM DTT, and 0.05% NP-40. After homogenization, samples were incubated on ice for 10 minutes and then spun at 3000 rpm for 10 minutes in a tabletop microfuge. Supernatant was spun at 14,000 x g for 10 minutes and supernatant was cytoplasmic fraction. Pellet from first spin was rinsed in PBS. Pellet was then resuspended in: 5mM HEPES, pH7.6, 1.5mM MgCl_2_, 300mM NaCl, 0.2mM EDTA, and 26% glycerol. Nuclei in resuspended pellet were lysed by 20 strokes with the large pestle of a Dounce homogenizer. Sample was incubated on ice for 30 minutes and then spun at 24,000 x g for 20 minutes. Supernatant was nuclear fraction. All buffers were pre-chilled on ice and contained protease inhibitors (Pierce). All spins were conducted at 4°C.

### PLA

Proximity ligation assay was performed using a Duolink In Situ Orange Mouse/Rabbit Kit (Sigma) according to the manufacturer’s instructions with the following modifications. Samples were dissected and fixed as for immunofluorescence. Tissue was permeabilized in PBST then washed 3x in PBS before proceeding. All subsequent steps were performed in recommended volumes in tubes with mounting of samples onto slides as the final step. Confocal images were taken on a Zeiss LSM 700 scanning microscope. For quantification, the total number of PLA puncta were counted in a maximum projection of a 101.61 x 101.61 x 24 µm volume from two regions of each disc: control and *dlg* RNAi. Presented images are a maximum projection of 5µm depth. Images were processed and analyzed in Fiji. Statistics were done in Prism.

## Acknowledgements

We thank Laura Mathies for cloning *Dlg*^*2XNLS>A*^, Norbert Perrimon for key reagents, the UC Davis Core Proteomics facility for assistance and the Bilder lab for helpful discussions. LC-MS was supported by a NIH shared instrumentation grant S10OD021801. This work was supported by NIH grants R01 GM090150 and R35 GM130388 to D.B., American Heart Association (AHA) Postdoctoral Fellowship 17POST33660155 to K.A.S. and AHA Predoctoral Fellowship 20PRE35120150 to M.J.K.

## SUPPLEMENTAL MATERIALS

**Supplemental Figure 1:**
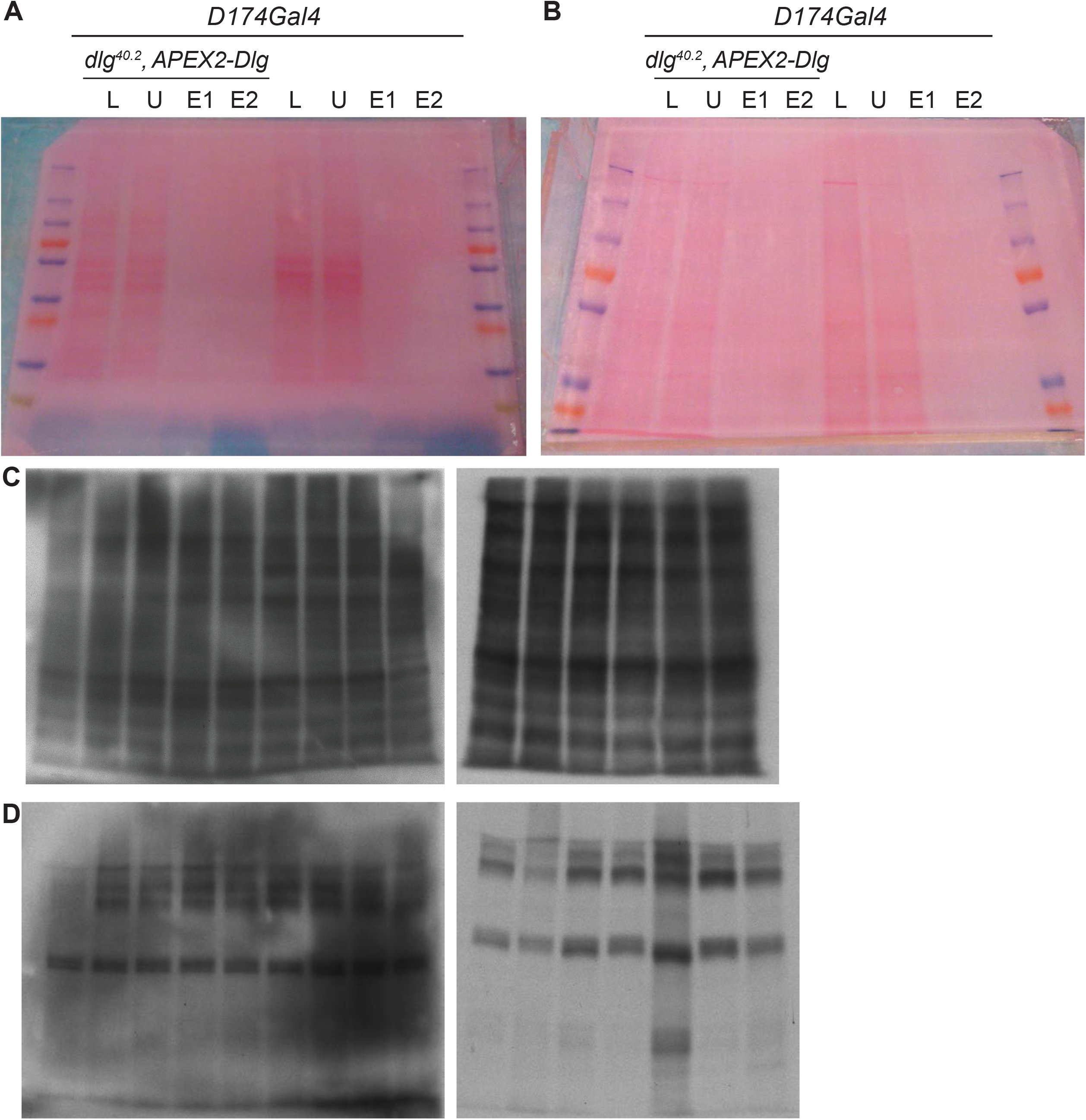
Loading and labeling controls. **A-B**. Ponceau-S staining as a protein loading control for Western blots in Figures 1G-H, showing equal amount of protein loaded in the lysate (L) and unbound (U) lanes for both experimental and control samples. Protein loading of eluates was below detection level of PonceauS staining, but efficient isolation of biotinylated proteins and subsequent elution from streptavidin beads was seen (Figures 1G-H). **C-D**. Streptavidin-HRP signal from MS sample lysate batches after labeling reaction but before pull-down with streptavidin beads. Each experimental (**C**) and control (**D**) batch showed clear bands without signs of protein degradation. Experimental batches showed consistent biotin labeling catalyzed by APEX2.

**Supplemental Figure 2:**
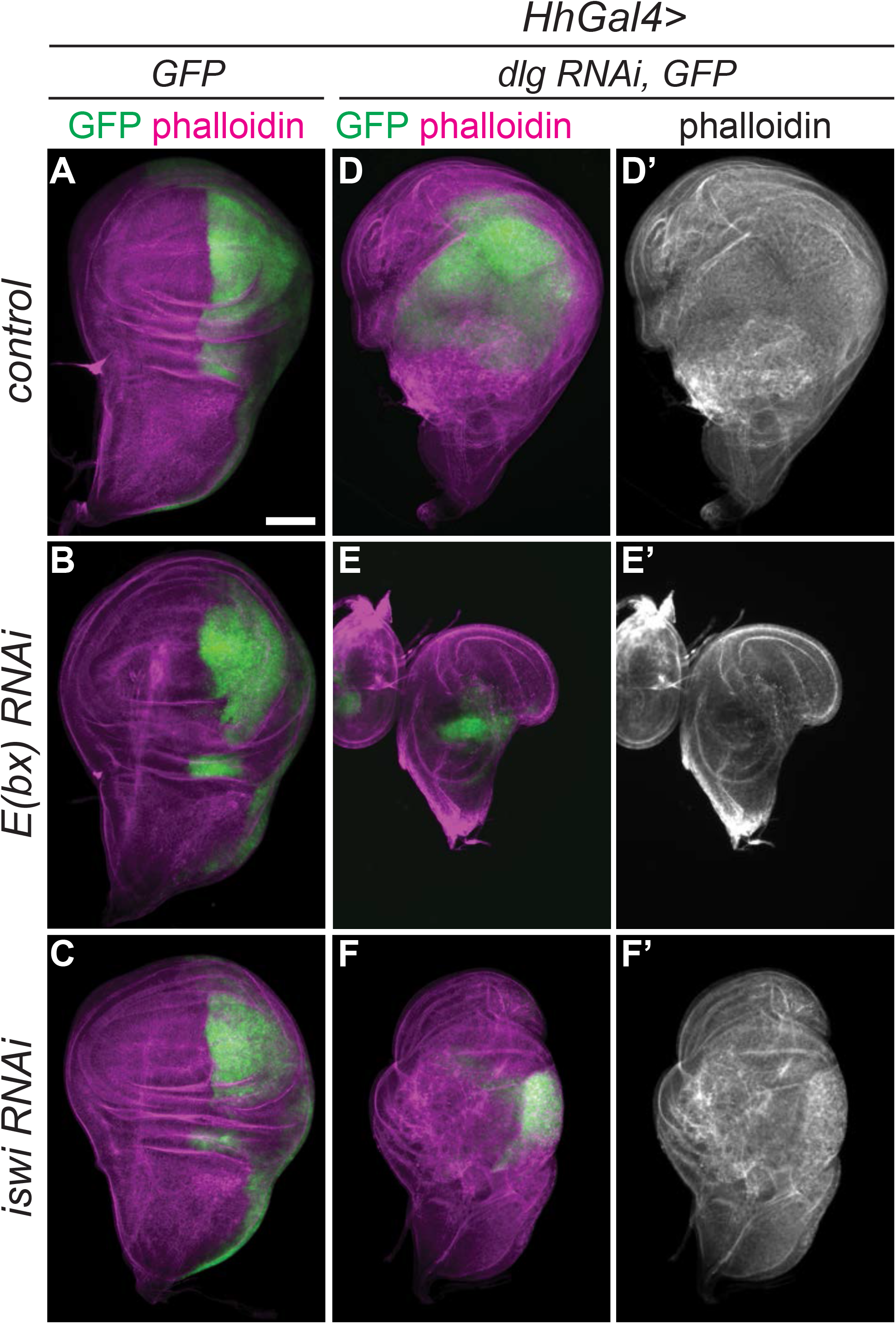
NURF RNAi reduces overgrowth in other *dlg* RNAi tumor systems. **A-C**. *hhGAL4* drives expression of *UAS-GFP* in the posterior compartment of wing discs (**A**). This expression domain and disc morphology were unaffected by either *E(bx)* (**B**) or *iswi* (**C**) RNAi. **D-F**. Expressing *dlg* RNAi under *hhGAL4* control causes overgrown, neoplastic tumors (**D**). Co-expression of *E(bx)* (**E**) or *iswi* (**F**) RNAi reduced the size of these tumors but did not rescue tissue architecture. **G-I**. Control eye disc size and morphology (**G**) were not affected by the expression of either *E(bx)* (**H**) or *iswi* (**I**) RNAi under the control of *eyFLP; FLPoutGAL4*.

**Supplemental Figure 3:**
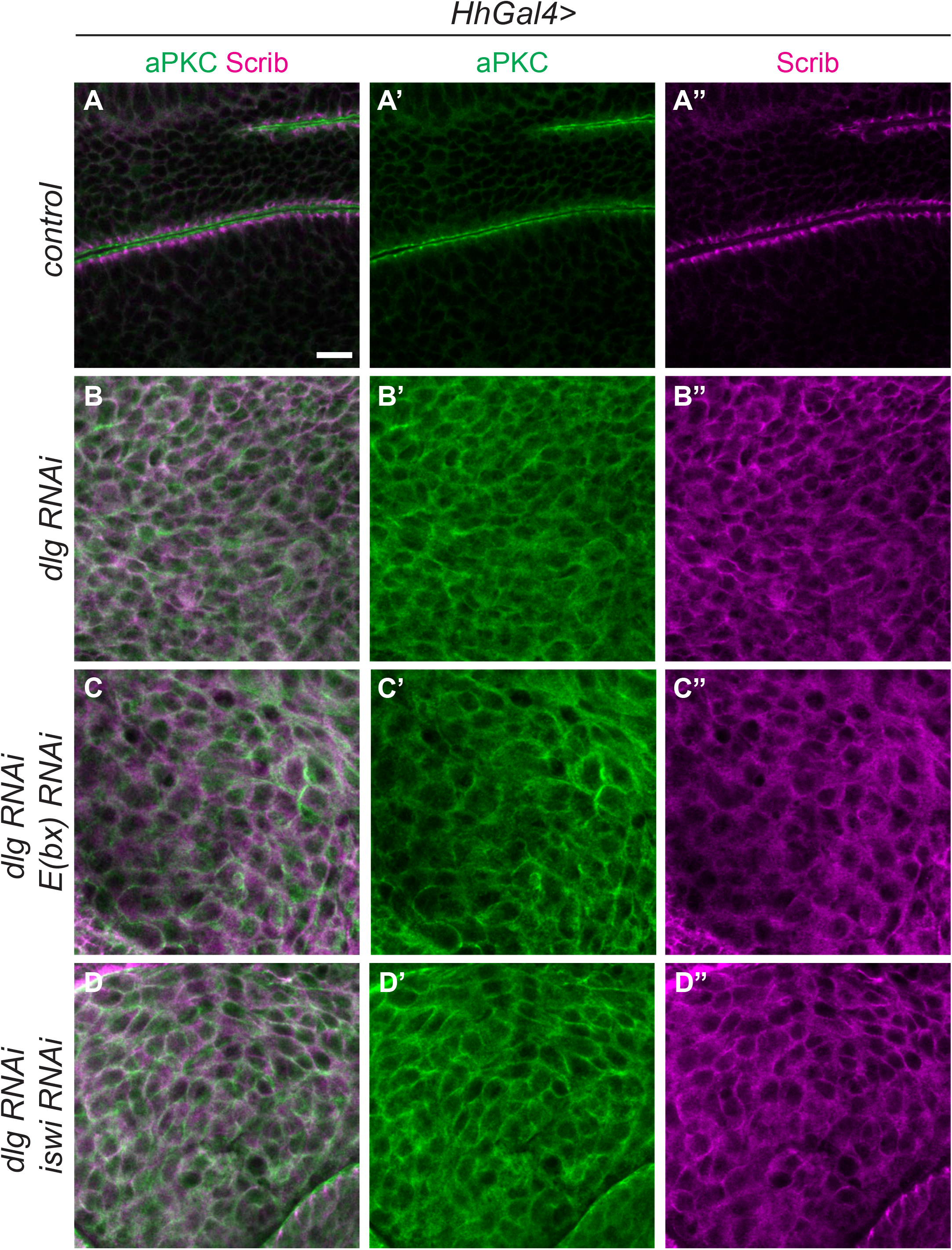
NURF RNAi does not rescue cell polarity in *dlg* RNAi tumors. **A-D**. Control wing disc cells (**A**) exhibit apicobasal polarity, as shown by apical localization of aPKC (**A’**) and basolateral localization of Scribble (**A”**). *dlg*-depleted cells (**B**) show loss of polarity and mixing of apical (**B’**) and basolateral (**B”**) markers. Polarity loss in *dlg* RNAi cells is not rescued by co-depletion of E(bx) (**C**) or Iswi (**D**). Scale bars, 10µm.

**Supplemental Figure 4:**
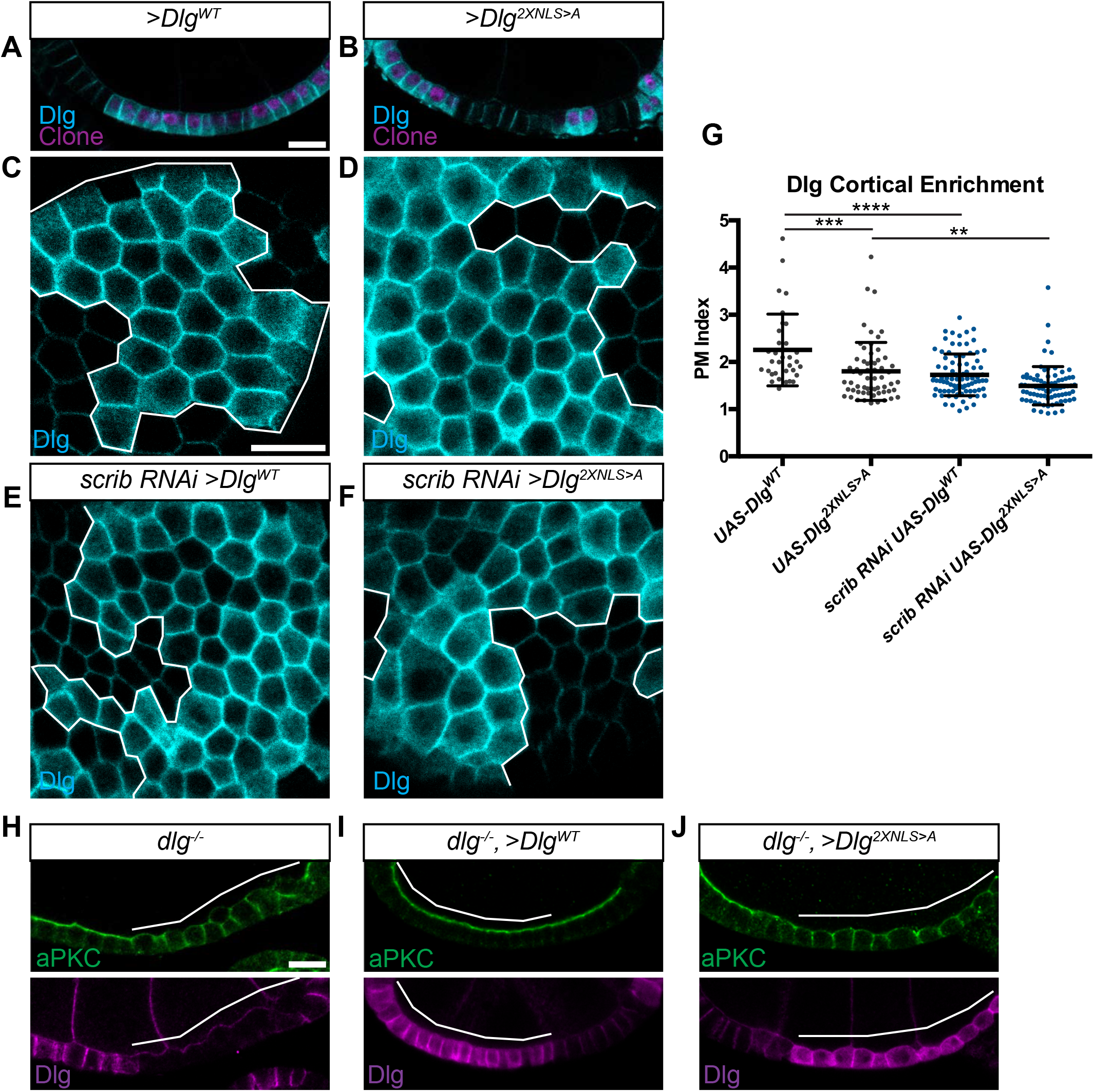
Transgenic Dlg localization in follicle epithelia. **A-E**. Clones overexpressing Dlg^WT^ (**A, C**) compared to Dlg^2XNLS>A^ (**B**,**D**) in cross-section (**A, B**) or en face (**C**,**D**) views. White lines indicate clone boundaries. Localization is similar, although Dlg^2XNLS>A^ shows 20% lower cortical enrichment (**G**). **E-F**. When expressed in a *scrib*-depleted background, both Dlg^WT^ (**E**) and Dlg^2XNLS>A^ (**F**) show reduced cortical enrichment than when expressed in a WT background (**G**) **H-J**. *dlg* null mutant follicle cell clones (**H**) are rescued by expression of Dlg^WT^ (**I**) but not Dlg^2XNLS>A^ (**J**). (**G**) One-way ANOVA with Tukey’s multiple comparisons test. Error bars represent S.D., data points represent measurements from single cells. **P<0.01, ***P<0.001, ****P<0.0001. Scale bars, 10µm.

**Supplemental Table 1: Complete Dlg-APEX2 proteomics results**

**Supplemental Table 2: Curated Dlg-APEX2 proteomics hit list**

**Supplemental Table 3: Dlg-APEX2 proteomics GO analysis**

**Supplemental Table 4: Genotypes, antibodies, chemicals and software used**

